# Alpha-180 spin-echo based line-scanning method for high resolution laminar-specific fMRI

**DOI:** 10.1101/2023.05.09.540065

**Authors:** Sangcheon Choi, David Hike, Rolf Pohmann, Nikolai Avdievich, Lidia Gomez-Cid, Weitao Man, Klaus Scheffler, Xin Yu

**Author notes:** **Lead corresponding author:** Dr. Xin Yu, Address: 13^th^ Street, Charlestown, MA 02129, USA.

## Abstract

Laminar-specific functional magnetic resonance imaging (fMRI) has been widely used to study circuit-specific neuronal activity by mapping spatiotemporal fMRI response patterns across cortical layers. Hemodynamic responses reflect indirect neuronal activity given limit of spatial and temporal resolution. Previous gradient-echo based line-scanning fMRI (GELINE) method was proposed with high temporal (50 ms) and spatial (50 µm) resolution to better characterize the fMRI onset time across cortical layers by employing 2 saturation RF pulses. However, the imperfect RF saturation performance led to poor boundary definition of the reduced region of interest (ROI) and aliasing problems outside of the ROI. Here, we propose α (alpha)-180 spin-echo-based line-scanning fMRI (SELINE) method to resolve this issue by employing a refocusing 180° RF pulse perpendicular to the excitation slice. In contrast to GELINE signals peaked at the superficial layer, we detected varied peaks of laminar-specific BOLD signals across deeper cortical layers with the SELINE method, indicating the well-defined exclusion of the large drain-vein effect with the spin-echo sequence. Furthermore, we applied the SELINE method with 200 ms TR to sample the fast hemodynamic changes across cortical layers with a less draining vein effect. In summary, this SELINE method provides a novel acquisition scheme to identify microvascular-sensitive laminar-specific BOLD responses across cortical depth.

## INTRODUCTION

Line-scanning fMRI has been successfully applied to investigate circuit-specific neuronal activity by measuring dynamic hemodynamic responses across cortical layers with high spatiotemporal resolution^1–9^. This is initially originated from Mansfield’s line-profile mapping studies in early 1970s^10,11^. The advantage of the current line-scanning fMRI method is to sample cortical layers with ultra-high spatial resolution. Meanwhile, the line-scanning method only acquires a single k-space line per timepoint, enabling an ultrafast sampling rate. This high spatiotemporal laminar fMRI sampling scheme has been being utilized for bottom-up and top-down blood-oxygenation-level-dependent (BOLD) fMRI mappings in both animal and human fMRI studies. Previously, Yu et al. developed a line-scanning fMRI method to delineate laminar fMRI onset time with distinct laminar-specific neural inputs such as thalamocortical input and corticocortical input in the rat brain with high spatial (50 um) and temporal resolution (50 ms)^1^. Line-scanning fMRI has been also combined with optogenetic control to further investigate the temporal features of the fast neural inputs across cortical layers in rodents^2^. Beyond preclinical fMRI studies, line-scanning fMRI for human brain mapping demonstrated a good correspondence with BOLD responses of 2D echo planar imaging (EPI) at the same temporal scale (200 ms)^12^. This line-scanning fMRI also motivated the cortical depth-dependent diffusion-based fMRI mapping schemes^13^. Lately, the ultra-fast line-scanning fMRI with k-t space reshuffling scheme has even provoked some interesting investigation of direct neuronal activity measurements^14^.

Typical gradient echo (GRE)-based line-scanning fMRI (GELINE) method needs to dampen signals outside of the region of interest (ROI) to avoid aliasing artifacts along the phase encoding direction^1,2,4,5,7–9^. Two saturation slices with additional RF exposure are applied for this purpose. However, two issues should be further investigated. One is the imperfect elimination of the aliasing artifacts (including inflow effects) due to imperfect RF performance and inhomogeneous B0 field. The other is specific absorption rate (SAR) problem with high duty cycle sequences. Here, we developed α (alpha)-180 line-scanning fMRI method to solve these problems. We modified spin-echo (SE) sequence by altering the refocusing 180° RF pulse perpendicular to the excitation slice^3,10,11^. This adjustment allows to only highlight a line-profile across the cortical layers without additional saturation RF pulses. In contrast to the GELINE method, SE based line-scanning fMRI (SELINE) method effectively exclude the surface draining vein effects. However, it should be noted that the laminar patterns of BOLD signals in SELINE can still be highly varied across different cortical layers in anesthetized rats. Furthermore, we also shorten TR to 200 ms for the SELINE method to sample the high resolution T2-weighted fMRI signals, demonstrating the feasibility of the fast sampling of laminar fMRI with effective ROI selectivity in rodents.

## RESULTS

### Mapping the evoked BOLD fMRI signals with GELINE and SELINE

We developed the SELINE method to map laminar-specific BOLD responses across cortical layers at the primary forepaw somatosensory cortex (FP-S1) of anesthetized rats, which can be compared with the conventional GELINE method^1^. First, unilateral electrical stimulation of left forepaw of rats showed robust BOLD responses in the right FP-S1 using EPI-fMRI method (**Fig. 1A**). Using the GELINE method, the selected FOV was defined by two saturation slices to avoid aliasing problem along the phase encoding direction (**Fig. 1B**). In contrast, the same FOV could be selected by applying a refocusing 180° RF pulse perpendicular to the excitation slice with SELINE (**Fig. 1E**). To compare ROI selectivity between GELINE and SELINE, 2D in-plane images were acquired by turning on phase encoding gradient (**Fig. 1C** and **1F**) and 1D profiles were plotted by averaging all readout voxels of the 2D image (**Fig. 1D and 1G**). Background signals were estimated from the areas outside of the FOV (for details, see the Method section): For GELINE, trial #1) 17.6 %, #2) 51.0 %, and for SELINE: trial #1) 2.3 %, #2) 3.0 %. This result indicated the efficiency of the SELINE method to produce sharper 2D slice profiles and lower background signals.

**Figure 1.**
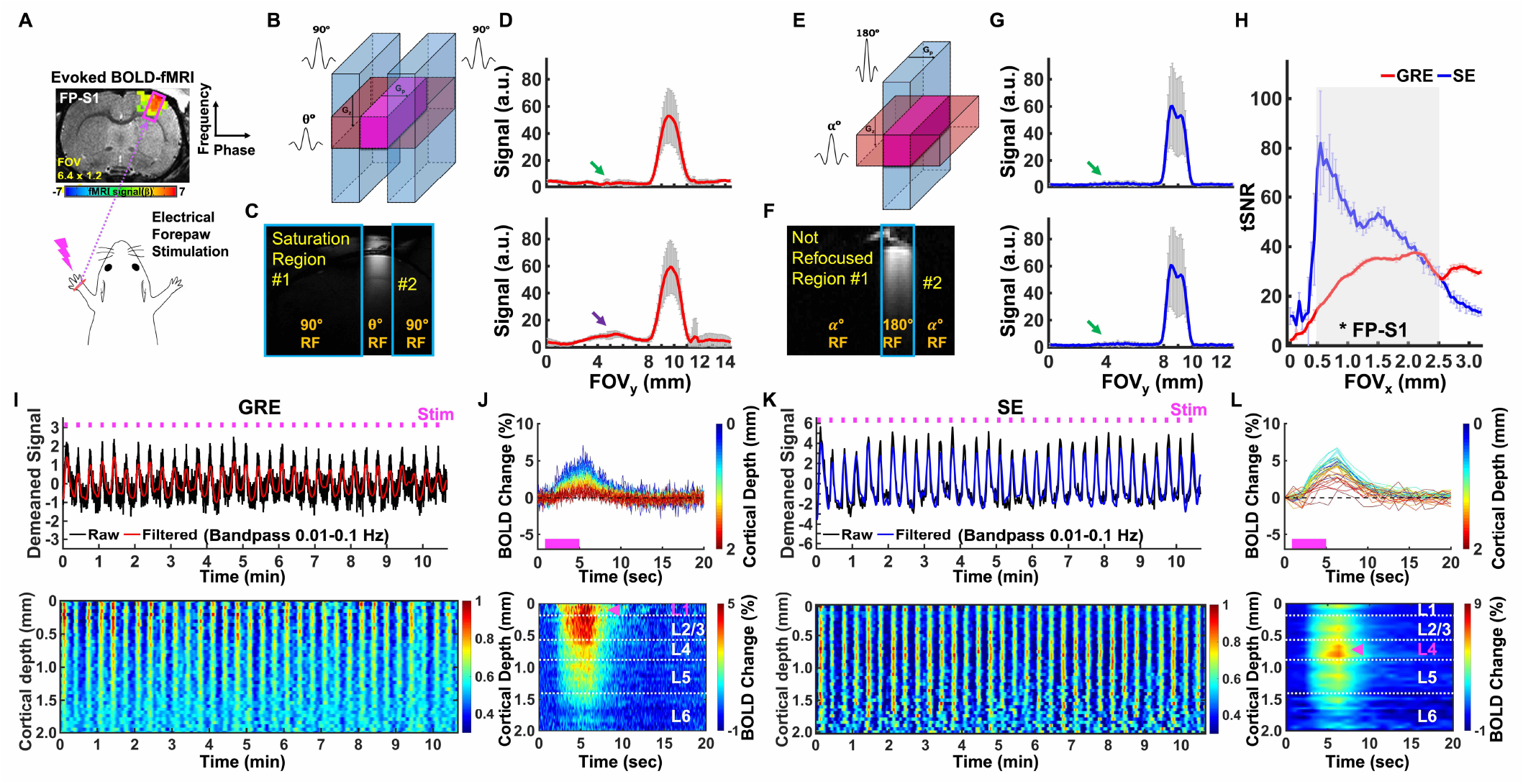
Evoked BOLD responses upon left forepaw stimulation using the GELINE and SELINE methods. **A**. Schematic illustration of the evoked fMRI experimental design on the EPI-BOLD activation map of FP-S1 region overlaid on an anatomical RARE image. **B-C**. Schematic drawing of GELINE imaging (**B**) and an acquired 2D image of GELINE (**C**). **D**. two representative 2D line-profiles of GELINE (average of 40 voxels): good saturation (green arrow) and bad saturation (purple arrow). Error bars represent mean ± SD across the cortical depths (0-2 mm). **E-F**. Schematic drawing of SELINE imaging (**E**) and an acquired 2D image of SELINE (**F**). **D**. two representative 2D line-profiles of SELINE (average of 40 voxels): good saturation (green arrows). Error bars represent mean ± SD across the cortical depths (0-2 mm). **H**. tSNR comparison between GELINE and SELINE (t-test: ^*^p < 10^−12^). **I-J**. A representative trial of GELINE. **I**. *Top*: Demeaned fMRI time series (32 epochs, 10 min 40 sec) of raw (black) and filtered (red) data (average of 40 voxels, bandpass: 0.01-0.1 Hz) in the FP-S1 region during electrical stimulation (3 Hz, 4 s, 2.5 mA) to left forepaw. *Bottom*: Normalized spatiotemporal map of the laminar-specific responses along the cortical depths (0–2 mm, 50 μm resolution). **J**. *Top*: Average BOLD time courses and *Bottom*: Average percentage change map across the cortical depths (0-2 mm, 40 lines in total) in the FP-S1. **K-L**. A representative trial of SELINE. **K**. *Top*: Demeaned fMRI time series (32 epochs, 10 min 40 sec) of raw (black) and filtered (red) data (average of 40 voxels, bandpass: 0.01-0.1 Hz) in the FP-S1 region during electrical stimulation (3 Hz, 4 s, 2.5 mA) to left forepaw. *Bottom*: Normalized spatiotemporal map of the laminar-specific responses along the cortical depths (0–2 mm, 50 μm resolution). **L**. *Top*: Average BOLD time courses and *Bottom*: Average percentage change map across the cortical depths (0-2 mm, 40 lines in total) in the FP-S1. Pink arrows indicate peak BOLD signals across the cortical layers.

To study the laminar fMRI characteristics of GELINE and SELINE across the cortical layers, we calculated temporal signal-to-noise ratio (tSNR) with 1D line-profiles which were acquired by turning off the phase encoding gradient. The tSNR of SELINE was higher than those of GELINE (**Fig. 1H**). The tSNR graph of SELINE had gradually decreasing trend across the cortical depth while those of GELINE had gradually increasing trend. The difference was likely caused by different TRs (1000 ms vs. 100 ms) and flip angles (90° vs. 50°) of the transceiver surface coil.

As shown in **Fig. 1I-L**, we demonstrated dynamic BOLD responses across different cortical layers of FP-S1 from the representative trial in individual GELINE (**Fig. 1I** and **1J**) and SELINE (**Fig. 1K** and **1L**) studies. **Fig. 1I** demonstrated periodic evoked BOLD signals upon left forepaw electrical stimulation with the T2*-weighted GELINE method, showing the dynamic laminar-specific BOLD responses as a function of time peaked around the superficial layer in the FP-S1 (4 s on/16 s off for each 20 s epoch, total 32 epochs). Average BOLD time series and laminar-specific BOLD maps illustrated that the peak BOLD response is located at L1, highlighting large draining vein effects at the cortical surface^15–21^ (**Fig. 1J**). In comparison to GELINE, SELINE also detected robust FP-S1 BOLD signals across different cortical layers (**Fig. 1K**), but showed peak BOLD signal located at L4, presenting improved spatial specificity to deeper cortical layers^15–18,20,21^ (**Fig. 1L**).

### Comparison of the laminar-specific peak BOLD responses in GELINE and SELINE

We further investigated the reproducibility of laminar-specific peak BOLD responses, as well as the variability of laminar-specific BOLD response patterns, between two methods (14 trials from 3 animals). The GELINE method detected peak BOLD signals primarily located at L1, but the peak BOLD signal detected by the SELINE method is much deeper. In animal #3, the ultra-strong BOLD signal detected in the superficial voxel indicates a large draining vein dominating the voxel BOLD signal. A similar BOLD response was also detected by the SELINE method, which may be contributed by potential intravascular effect of which the large draining vein is not negligible in the voxels with only 50 um thickness (**Fig. 2G** and **2H**). Interestingly, the layer-specific BOLD signal varies largely across animals in both GELINE and SELINE maps. Besides the primary peak BOLD in L1 of GELINE, a second peak appeared in L4 in some animal (**Fig. 2D**). And for SELINE method, the primary peak also varies at L2/3 and L4, which present highly different laminar patterns from GELINE even acquired from the same animal with interleaved trials during experiments. These results have suggested that the profile of laminar-specific BOLD signals can vary largely across animals, which may present varied dynamic patterns of BOLD responses due to the altered neurovascular coupling across different cortical layers.

**Figure 2.**
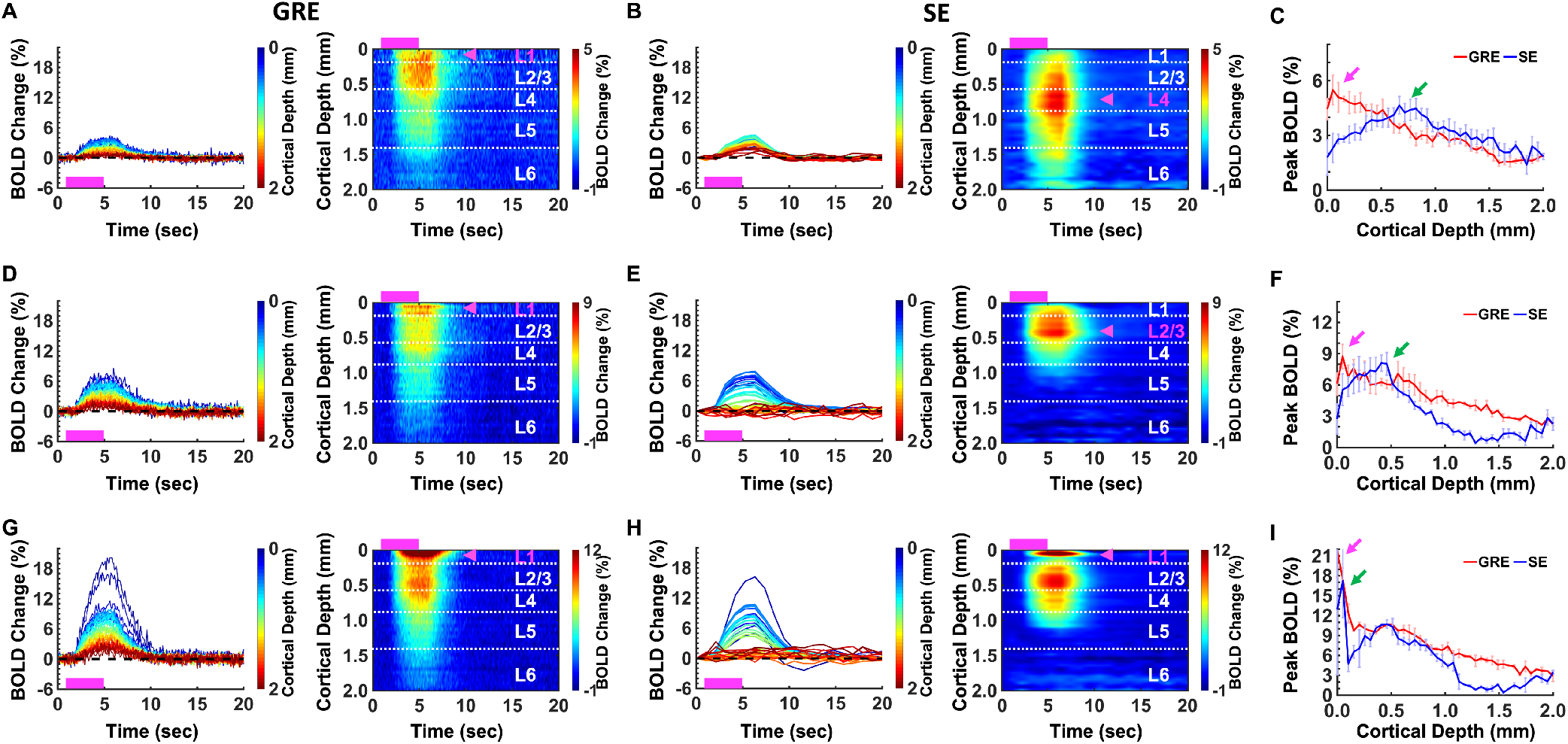
Evoked fMRI time series and percentage change maps of GELINE and SELINE in rat brains (14 trials of 3 rats). **A-C**. Rat #1 (3 trials of each). **D-F**. Rat #2 (2 trials of each). **G-I**. Rat #3 (2 trials of each). **A, D, G**. *Left*: Average BOLD time courses and *Bottom*: Average percentage change map of GELINE across the cortical depths (0-2 mm, 40 lines in total) in FP-S1 region. **B, E, H**. *Left*: Average BOLD time courses and *Bottom*: Average percentage change map of SELINE across the cortical depths (0-2 mm, 40 lines in total) in FP-S1 region. Pink boxes indicate stimulation duration and pink arrows indicate peak BOLD signals across the cortical layers. **C, F, I**. Comparison of peak BOLD signals between GELINE (pink arrows) and SELINE (green arrows). Error bars represent mean ± SD of peak BOLD signals.

### Mapping the laminar BOLD responses with a 200 ms SELINE method

We performed BOLD fMRI experiments with 200 ms time of repetition (TR) by applying optimized flip angles based on the Bloch equation^22,23^. For comparison, we also performed GELINE method in the same anesthetized rat. As shown in **Fig. 3A-D**, we demonstrated the evoked BOLD responses across the cortical layers upon the periodic electric stimulation with the GELINE (**Fig. 3A** and **3B**) and SELINE (**Fig. 3C** and **3D**) methods, showing the average BOLD time series and percentage changes peaked at L1 in both GELINE and SELINE. To characterize the laminar-specific BOLD responses, the normalized BOLD signals were plotted across the cortical layers. As shown in **Fig. 3F**, the GELINE method had the steep signal drop from L1 to L2, while the SELINE method had the gradual signal drop across the cortical depth. It indicates that the high temporal SELINE method reduces the large vessel contribution to the BOLD responses by minimizing magnetic susceptibility effects at the superficial layer (i.e., L1). To select an optimized flip angle, the tSNR of different flip angles was plotted (**Fig. 3G**). Even though the optimal flip angle for TR 200 ms was ∼150° and had the highest tSNR, the difference of the tSNR change was relatively small in multiple trials with the different flip angles. This result was possibly caused by the long T1 effect (∼2200 ms) in SELINE acquisition with a short TR (200 ms)^24^. As same as the theoretical predictions based on the Bloch equation ^22,23^, *e*^-*TR*/*T*1^ was almost close to one and thus, the maximum intensity at the optimal flip angle wasn’t changed much. It was noteworthy that the average tSNR of SELINE was higher at the superficial and middle layers than that of GELINE (**Fig. 3E** and **3G**) due to larger flip angle (100-150° vs. 50°) and longer TR (200 ms vs. 100 ms). In summary, these results not only demonstrated less magnetic susceptibility effects at the superficial layer, but also highlighted both tSNR and laminar specificity enhancement in SELINE with high temporal resolution.

**Figure 3.**
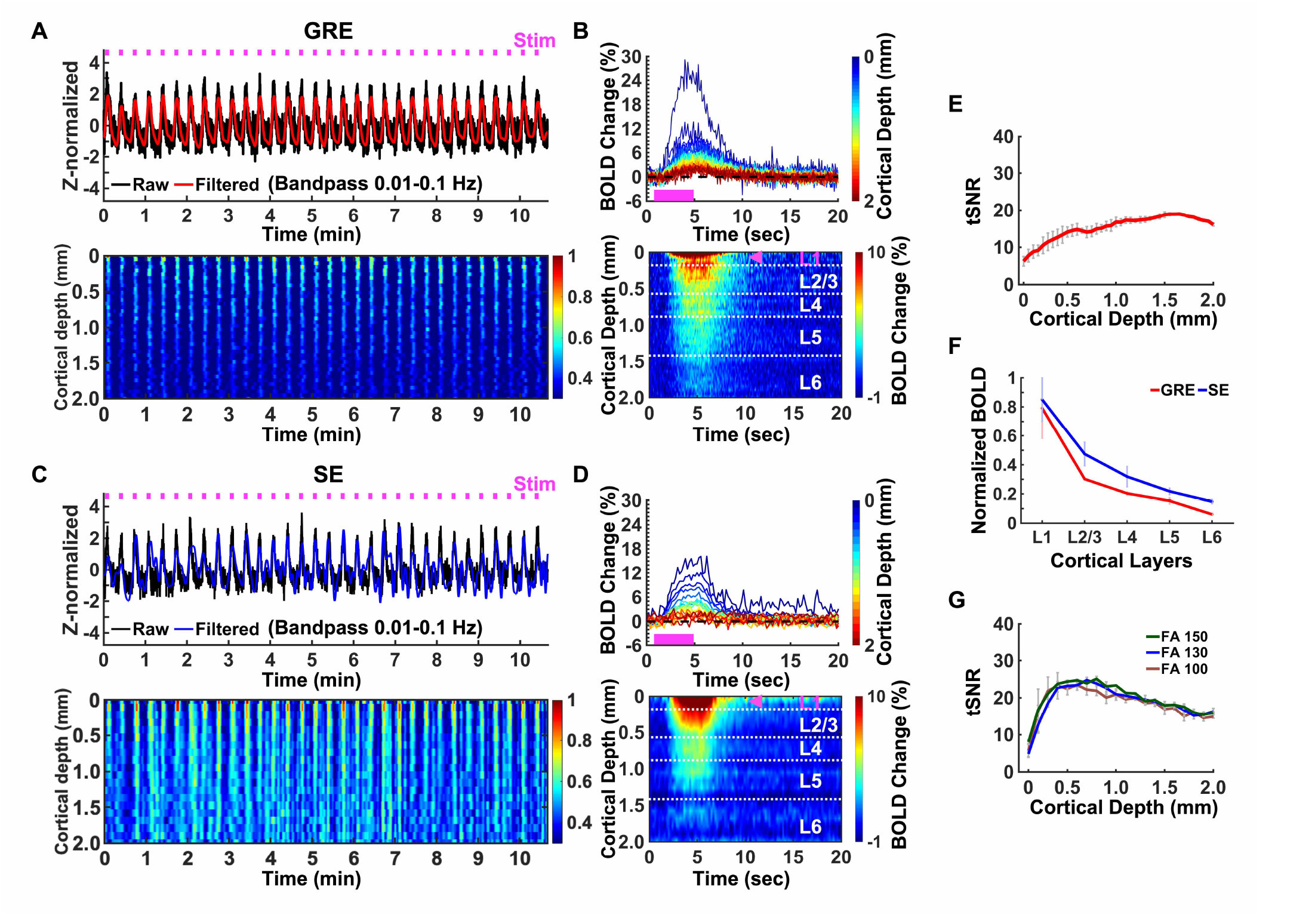
Evoked fMRI responses with GRE (TR 100 ms) vs. SE (TR 200 ms). **A-B**. GELINE (2 trials) **A**. *Top*: Z-score normalized fMRI time series (average of 40 voxels) of FP-S1. *Bottom:* Normalized spatiotemporal map of the laminar-specific responses along the cortical depths (0–2 mm, 50 μm resolution). **B**. *Top*: Average BOLD time courses and *Bottom*: Average percentage change map across the cortical depths (0-2 mm, 40 lines in total) in the FP-S1. **C-D**. SELINE (3 trials) **C**. *Top*: Z-score normalized fMRI time series (average of 20 voxels) of FP-S1. *Bottom*: Normalized spatiotemporal map of the laminar-specific responses along the cortical depths (0–2 mm, 100 μm resolution). **D**. *Top*: Average BOLD time courses and *Bottom*: Average percentage change map across the cortical depths (0-2 mm, 20 lines in total) in the FP-S1. **E**. tSNR of GELINE (2 trials) across the cortical depths (0-2 mm). **F**. Comparison of normalized BOLD signals between GELINE and SELINE across cortical layers. **G**. tSNR comparison of SELINE with three excitation flip angles (calculated by the Bloch simulation) across the cortical depths (0-2 mm): FA 100 (5 trials), FA 130 (3 trials), and FA 150 (3 trials). Pink boxes indicate stimulation duration and pink arrows indicate peak BOLD signals across the cortical layers. Error bars represent mean ± SD.

## DISCUSSION

In this study, we applied the SELINE method to investigate laminar-specific evoked BOLD responses across cortical layers with high spatial and temporal resolution. The SELINE method has sharper and better ROI-selectivity than the GELINE method, employing the refocusing 180 RF pulse perpendicular to the excitation plane. Our results show that the peak signal of SELINE is spread across the cortical layers while those of GELINE is at the superficial layer^25,26^By pushing the temporal resolution of SELINE to 200 ms, we also demonstrate the feasibility to map laminar-specific BOLD response with the suppression of the large draining vein effect^15–18,20,21^ in comparison to the GELINE method.

Significant effort with high magnetic field fMRI has been made to explore laminar fMRI responses corresponding to distinct information flows (e.g., top-down/bottom-up or feedforward/feedback) at high spatial and temporal scales in both animals and humans. Among these efforts, cortical depth-dependent fMRI, detecting BOLD, cerebral blood volume (CBV), and cerebral blood flow (CBF) signals with both SE and GRE methods, has identified hemodynamic regulation, blood volume distribution, circuit-specific laminar responses, and hierarchical information streams across cortical layers in animal^1,2,5,8,15,16,25–29^ and human brains^30–34^. In particular, the high resolution CBV-fMRI (based on the VASO mapping scheme) has been used to measure layer-specific directional functional connectivity across human motor cortex and somatosensory and premotor regions^30^. It should be noted that the cortical thickness of human brains is in the range of 1-4 mm, which is highly comparable to that of rodent brains in the range of 1-2 mm^35^. Given the limited spatial resolution of the high field laminar-fMRI method (∼600-700 um), the truly counted voxels across different cortical regions are in the single digit number, which could be much better improved by the developed line-scanning fMRI method, as well as with ultra-fast sampling rate.

Recently, the GRE-based line-scanning BOLD mapping scheme has been implemented to investigate BOLD signals across cortical layers in human fMRI studies^7,12^. Nevertheless, the required saturation RF pulses of the GELINE method result in high specific absorption rate (SAR) and total RF power limits with short TRs, inducing more complicated aliasing problems. For the SELINE method, the beam-like line-scan projection has been previously applied for probing myeloarchitecture across cortical layers in the primary somatosensory cortex (S1) and primary motor cortex (M1) of the human brain^13^ and mapping irreversible and reversible transverse relaxation rates (i.e., R2 and R2′) in primary visual cortex (V1), S1, and M1 of human brains^36^. We thus applied this SELINE method to better characterize layer-specific fMRI features across cortical depths at FP-S1 of rodent brains. The SELINE method employed the spin-echo scheme to reduce the large draining vein effect, which can be further distinguished from the deeper cortical layer responses given the high spatial resolution (**Fig. 1I-L**).

As reported in previous studies^15–17,25,37–39^, GELINE is more sensitive to large veins at the pial surface and has poor specificity across different cortical depths, whereas SELINE is less vulnerable to superficial large draining veins and has good sensitivity to microvessel across cortical layers. However, the largely varied laminar patterns of the BOLD responses were observed in both methods (**Fig. 2**). It may suggest that the varied patterns of laminar-specific BOLD signals pertain on microvascular biases and baseline blood volume distribution across cortical layers ^40,41^. Whereas the confounding observation of the varied peak profiles of BOLD responses across different cortical layers, these results illustrate the feasibility of the line-scanning method to detect distinct laminar BOLD responses, providing a high-resolution mapping scheme when investigating altered neurovascular coupling events across cortical layers.

The limitation of SELINE is the slow sampling rate. We tried to shorten TR by adjusting excitation flip angle. Based on Bloch equation^22,23^, we have estimated the appropriate angles with a short TR (i.e., 200 ms). Our results show the feasibility of the fast SELINE method which has a good sampling capability capturing dynamic BOLD signals from superficial to deeper layers. For the future work, simultaneous GRE- and SE-type fMRI acquisition can be applied to better characterize laminar-specific fMRI patterns and minimize time dependency of dynamic fMRI responses by employing GRASE^42^-based line-scanning in rodents as already suggested for the human fMRI mapping^6^.

## METHODS

### Animal preparation

The study was performed in accordance with the German Animal Welfare Act (TierSchG) and Animal Welfare Laboratory Animal Ordinance (TierSchVersV). This is in full compliance with the guidelines of the EU Directive on the protection of animals used for scientific purposes (2010/63/EU) and the MGH Guide for the Care and Use of Laboratory Animals. The study was reviewed by the ethics commission (§15 TierSchG) and approved by the state authority (Regierungspräsidium, Tübingen, Baden-Württemberg, Germany) and the MGH Institutional Animal Care and Use Committee (Charlestown, MA, USA). A 12-12 hour on/off lighting cycle was maintained to assure undisturbed circadian rhythm. Food and water were available ad libitum. A total of 4 male Sprague–Dawley rats were used in this study.

Anesthesia was first induced in the animal with 5% isoflurane in the chamber. The anesthetized rat was intubated using a tracheal tube and a mechanical ventilator (SAR-830, CWE, USA) was used to ventilate animals throughout the whole experiment. Femoral arterial and venous catheterization was performed with polyethylene tubing for blood sampling, drug administration, and constant blood pressure measurements. After the surgery, isoflurane was switched off, and a bolus of the anesthetic alpha-chloralose (80 mg/kg) was infused intravenously. After the animal was transferred to the MRI scanner, a mixture of alpha-chloralose (26.5 mg/kg/h) and pancuronium (2 mg/kg/h) was constantly infused to maintain the anesthesia and reduce motion artifacts.

### EPI fMRI acquisition

All data sets from rats were acquired using a 14.1T/26 cm (Magnex, Oxford) horizontal bore magnet with an Avance III console (Bruker, Ettlingen) and a 12 cm diameter gradient system (100 G/cm, 150μs rising time). A home-made transceiver surface coil with a 10 mm diameter was used on the rat brain. For the functional map of BOLD activation (**Fig. 1A)**, a 3D gradient-echo EPI sequence was acquired with the following parameters: TR/TE 1500/11.5 ms, FOV 1.92 × 1.92 × 1.92 cm^3^, matrix size 48 × 48 × 48, spatial resolution 0.4 × 0.4 × 0.4 mm^3^. A high order (*e*.*g*., 2^nd^ or 3^rd^ order) shimming was applied to reduce the main magnetic field (B0) inhomogeneities at the region-of-interest. For anatomical reference of the activated BOLD map, a RARE sequence was applied to acquire 48 coronal images with the same geometry as that of the EPI images. The fMRI design paradigm for each trial comprised 200 dummy scans to reach steady-state, 10 pre-stimulation scans, 3 scans during stimulation, and 12 post-stimulation scans with a total of 8 epochs.

### GELINE acquisition

GELINE datasets (9 trials of 4 rats) were acquired with a 6-mm diameter home-made transceiver surface coil in anesthetized rats for evoked fMRI. GELINE was applied by using two saturation slices to avoid aliasing artifacts in the reduced field-of-view along the phase encoding (*i*.*e*., from left to right) direction (**Fig. 1B** and **1C**). 2D line profiles were acquired to evaluate saturation RF pulses performance (**Fig. 1D**). Laminar fMRI responses were acquired along the frequency-encoding direction (**Fig. 1I** and **1J**). The following acquisition parameters were used: TR/TE 100/12.5 ms, TA 10 min 40 sec, FA 50°, slice thickness 1.2 mm, FOV 6.4 × 3.2 mm^2^, and matrix 128 × 32. The fMRI design paradigm for each epoch consisted of 1 second pre-stimulation, 4 seconds stimulation, and 15 seconds post-stimulation with a total of 20 seconds. A total of 6400 lines (*i*.*e*., 10 m 40 s) in each cortex were acquired every single trial in evoked fMRI. Evoked BOLD activation was identified by performing electrical stimulation to the left forepaw (300 µs duration at 2.5 mA repeated at 3 Hz for 4 seconds).

### SELINE acquisition

SELINE datasets (18 trials of 4 rats) were acquired in anesthetized rats for evoked fMRI. SELINE was applied by the 180° RF pulse oriented perpendicular to the α° excitation RF pulse as moving the refocusing gradient to phase encoding gradient in order to obtain high spatial resolution without reduced FOV aliasing problem along the phase encoding (*i*.*e*., from left to right) direction (**Fig. 1E** and **1F**). 2D line profiles were also acquired to evaluate the refocusing RF pulses performance (**Fig. 1G**). Laminar fMRI responses were acquired along the frequency-encoding direction (**Fig. 1K** and **1L**). The following acquisition parameters were used: TR/TE/FA 1000/20 ms/90°, 200/10 ms/ 100° or 130° or 150°, TA 10 min 40 sec, slice thickness 1.2 mm, FOV 3.2 × 1.2 mm^2^ for TR 1000 ms, FOV 6.4 × 1.2 mm^2^ for TR 200 ms, and matrix 64 × 32. The fMRI experiment set-up was identical to those of the GELINE in evoked fMRI.

### Data Analysis

All signal processing and analyses were implemented in MATLAB software (Mathworks, Natick, MA) and Analysis of Functional NeuroImages software^43^ (AFNI, NIH, USA). For evoked fMRI analysis for **Fig. 1A**, the hemodynamic response function (HRF) used was the default of the block function of the linear program 3dDeconvolve in AFNI. BLOCK (L, 1) computes a convolution of a square wave of duration L and makes a peak amplitude of block response = 1, with *g*(*t*) = *t*^4^*e*^-*t*^/[4^4^*e*^−4^]. Each beta weight represents the peak height of the corresponding BLOCK curve for that class. The HRF model was defined as follows:

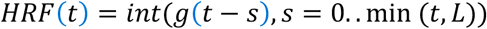

Cortical surfaces were determined based on signal intensities of fMRI line profiles as described in the previous work. The detailed processing was conducted as provided in the previous line-scanning studies ^1,5,8^. For Fig. 1I and 1K, demeaned fMRI time courses were used as follows: (x - μ), where x was the original fMRI time courses and μ was the mean of the time courses. The line profile map concatenated with the multiple fMRI signals was normalized by a maximum intensity. The Z-score normalized time courses were calculated as follows: (x - μ)/σ, where x was original fMRI time courses and μ, σ were the mean and the standard deviation of the time courses, respectively (zscore function in MATLAB). Average BOLD time series and percentage changes were defined as (S-S0)/S0 × 100 %, where S was the BOLD signal and S0 was the baseline. S0 was obtained by averaging the fluctuation signal in the 1-second pre-stimulation window in evoked fMRI that was repeated every 20 seconds with the whole time series (640 sec). The BOLD time series in each ROI were detrended (‘polyfit’ function in Matlab, order: 3) and bandpass filtered (0.01-0.1 Hz, FIR filter, order: 4096). The bandpass filtering was performed as a zero-phase filter by ‘fir1’ and ‘filter’ functions in Matlab, compensating a group delay (‘grpdelay’ and ‘circshift’ functions in Matlab) introduced by the FIR filter. Temporal signal-to-noise ratio (tSNR) values were calculated across the cortical depths to compare tSNR differences between GELINE and SELINE. Student t-test was performed with the tSNR values of GELINE and SELINSE (**Fig. 1H)**. The p-values < 0.05 were considered statistically significant.

### Bloch stimulation

To optimize the α° excitation flip angle with short TR (i.e., 200 ms) in SELINE, signal intensities were calculated as a function of excitation flip angle by solving the Bloch equation^22,23^, by employing the refocusing 180° RF pulse. The maximum signal intensity occurred at the Ernst angle which was defined as follow:

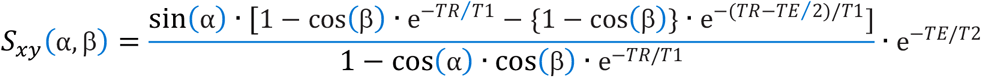

where α, β indicate excitation and refocusing flip angles, respectively and T1, T2 indicate longitudinal and transverse magnetization parameters, respectively. T1 and T2 values were estimated from the previous study^24^.

## Data availability

All other data generated during this study are available from the corresponding author upon reasonable request.

## Code availability

The related image processing codes are available from the corresponding author upon reasonable request.

## ACKNOWLEDGEMENTS

This research was funded by NIH funding (RF1NS113278, R01NS124778, R01NS122904, R01NS120594, R21NS121642), NSF grant 2123971, and the S10 instrument grant (S10 MH124733–01) to Martino’s Center, German Research Foundation (DFG) Yu215/2-1, 3-1, BMBF 01GQ1702, and the internal funding from Max Planck Society. We thank Dr. J. Engelmann, and Ms. H. Schulz for technical support, Dr. P. Douay, Ms. R. König, and Ms.M. Pitscheider for animal support, the AFNI team for the software support.

